# Phenotypic Heterogeneity Shapes Phage Resistance and Cocktail Efficacy in *Klebsiella pneumoniae*

**DOI:** 10.1101/2025.11.11.687788

**Authors:** Lucas Mora-Quilis, Rafael Sanjuán, Pilar Domingo-Calap

## Abstract

The emergence of phage-resistant bacteria poses a significant challenge to the success of phage therapy. Although phage cocktails can delay resistance, their efficacy relies on the ability to target the full spectrum of resistant variants, which are often more diverse than that represented by clonal isolates. In this study, we investigated how phenotypic diversity within phage-resistant *Klebsiella pneumoniae* influences susceptibility to newly isolated phages and the effectiveness of phage cocktails. We isolated phages from three cultures resistant to a capsule-dependent phage: a heterogeneous acapsular population, an acapsular mutant with a stable phenotype, and a capsule-reverted isolate that regained capsule expression upon removal of phage pressure. These phages differed in host tropism and in their capacity to delay resistance emergence when combined with the capsule-dependent phage. Phages targeting acapsular variants, particularly those isolated from the heterogeneous population, were the most effective, exhibiting strong synergy. Single-cell analyses further revealed that sustained selective pressure from the capsule-dependent phage prevents capsule reversion and maintains cocktail efficacy. Overall, our results highlight the importance of accounting for phenotypic heterogeneity when designing phage therapies and support population-level approaches for optimizing phage cocktail composition.

**Importance:** Phage therapy is a promising alternative to antibiotics, but its success is often limited by the rapid emergence of phage-resistant bacteria. These resistant populations can be highly heterogeneous, comprising both stable mutants and variants with reversible, non-genetic resistance. In this study, we explore how this phenotypic diversity influences the effectiveness of phage cocktails. By isolating new phages and testing them in combination, we demonstrate that the selective pressure exerted by specific phages can prevent the reversion in transiently resistant variants, thereby sustaining treatment efficacy. Our findings highlight the need to consider not only the range of bacterial targets but also how phage pressure shapes bacterial population dynamics. This work offers a more refined strategy for designing phage cocktails with improved clinical potential.

## INTRODUCTION

Phage therapy has re-emerged in recent years as a promising alternative or complement to traditional antibiotic treatments in the fight against multidrug-resistant bacteria (1, 2). Numerous clinical cases have been reported, and the number of clinical trials involving phage treatments has grown significantly in the last years (3, 4). However, the rapid emergence of phage-resistant bacteria remains one challenge limiting the efficacy of phage therapy (5–8). To address this issue, phage cocktails, combinations of two or more phages administered simultaneously, have been proposed as a strategy to both broaden host range and suppress resistance. Bacteria possess a vast and diverse array of mechanisms to evade phage infection that include adaptive immunity (CRISPR-Cas) (9), innate defence such as restriction-modification systems (10), programmed cell death through the abortive infection system (11), and many others (12, 13). However, one of the simplest and most effective strategies to prevent phage infection, is the modification of cell surface structures that phages use for host recognition. In *Klebsiella* spp., an enterobacteria with high antibiotic resistance rates, capsule is the primary factor determining phage infectivity (14, 15). Consequently, mutations that lead to capsule loss or receptor modification can confer resistance to capsule-dependent phages, with the trade-off of exposing other surface structures, such as lipopolysaccharides (LPS) or outer membrane proteins (OMPs), that may serve as receptor for other phages (16–19). Importantly, capsule expression is not governed by a single gene, but is influenced by a large set of genetic and regulatory factors, leading to a substantial variability in capsule quantity and architecture (20, 21). Similarly, LPS and OMP can also vary among cells within the population (22, 23), further diversifying susceptibility to non-capsule dependent phages. Modification of these bacterial structures can also result from reversible changes, such as phase variation or stochastic fluctuations in gene expression (24–27) (ref). These changes are highly dynamic and allow bacteria to rapidly switch between phenotypes, conferring a rapid adaptive strategy in changing environments, highlighting the potential adaptive advantage of transient resistance over mutation-based mechanisms. Interestingly, acapsular variants lacking the protective capsule have been shown to be less virulent, making them more susceptible to conventional antibiotics and clearance by the host immune system (28, 29). However, in immunocompromised patients or when antibiotics are no longer effective, these variants can spread rapidly and accelerate the onset of disease (30), requiring novel strategies of control. Indeed, phage cocktails composed primarily of phages targeting clonal resistant cultures have been proposed as a strategy to delay the emergence of phage resistance in *K. pneumoniae* (31–33). Nevertheless, the intrinsic heterogeneity of bacterial populations, including differential expression of the capsule components, may pose a challenge, making it difficult to capture the resistance diversity through the use of a single or few resistant clones for phage isolation (34).

In a previous study using *K. pneumoniae* capsule type 1 as a model, we showed that resistance to the capsule-dependent phage Cap62 resulted in a highly heterogeneous acapsular population (35). This population was composed predominantly of reversible acapsular variants, that transiently suppressed capsule expression through non-mutational mechanisms, and to a lesser extent, of stable acapsular mutants. While the mutants maintained an acapsular phenotype, the reversible variants rapidly restored capsule production once phage pressure was removed. This heterogeneity in the resistant populations poses a challenge for phage selection, as clonal isolates may not capture the full spectrum of resistance mechanisms. Therefore, here, we aimed to explore how phenotypic diversity in phage-resistant *K. pneumoniae* influences susceptibility to newly isolated phages with the potential to delay resistance emergence when used in cocktails. To this end, we isolated new phages using three Cap62-resistant cultures: the heterogeneous acapsular population, an acapsular mutant with stable phenotype, and a capsule-reverted isolate that regained capsule expression upon removal of phage pressure. The isolated phages exhibited different host tropism and different ability in delaying the emergence of resistance when combined with Cap62. Phages targeting acapsular variants were more effective in delaying resistance, particularly those isolated from the heterogeneous population, which exhibited a strong synergistic effect with Cap62. Finally, single-cell experiments demonstrated that the selective pressure exerted by Cap62 is essential to prevent capsule reversion and sustain the long-term efficacy of the phage cocktail.

## RESULTS

### Isolation and characterization of phages targeting the Cap62-resistant cultures

Cap62-resistant cultures were used as host for phage isolation. Phages CuaHET1 and CuaHET2 were obtained from the heterogeneous resistant population, including acapsular mutants and reversible acapsular variants. To preserve the acapsular phenotype during isolation, phage Cap62 was maintained as selective pressure, preventing capsule restoration and enriching for acapsular cells. In addition, phages CuaMUT1 and CuaMUT2 were isolated from the acapsular mutant, and phages CuaREV1 and CuaREV2 from the capsule-reverted isolate (Figure 1). The ability of these phages to infect the Cap62-resistant cultures was evaluated. Similar to phage Cap62, phages CuaREV1 and CuaREV2 were able to form spots and plaque-forming units (PFU) on both, the capsular WT strain and the capsule-reverted isolate. In contrast, phages CuaMUT1, CuaMUT2, CuaHET1 and CuaHET2 only were able to infect and lyse acapsular resistant cultures. Notably, these phages produced clear spots and PFUs only in the specific culture from which they were isolated (Figure 2A).

**Figure 1.**
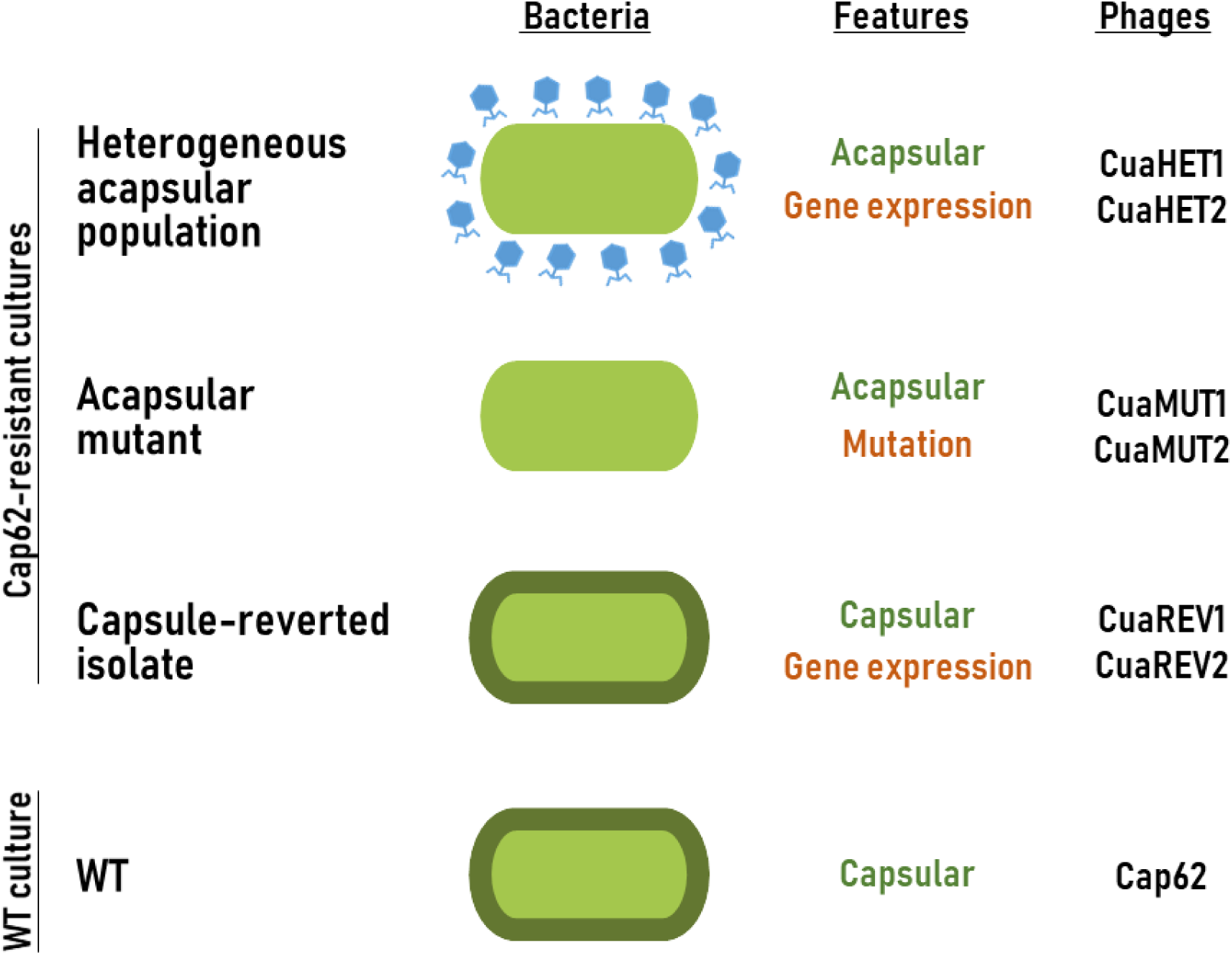
Bacterial cultures used for phage isolation. Illustrative scheme of the bacterial cultures used in this study (WT culture, and Cap62-resistant cultures: heterogeneous acapsular population, acapsular mutant and capsule-reverted isolate). Bacterial cells are depicted in green, with a dark green outline indicating the presence of capsule in the WT and the capsule-reverted isolate. The blue phage shown with the heterogeneous acapsular population represents phage Cap62, highlighting its essential role in maintaining the acapsular phenotype. Key features of each strain are indicated: The presence of capsule (green) and the type of resistance (orange). In addition, the specific phages isolated in each strain are listed.

**Figure 2.**
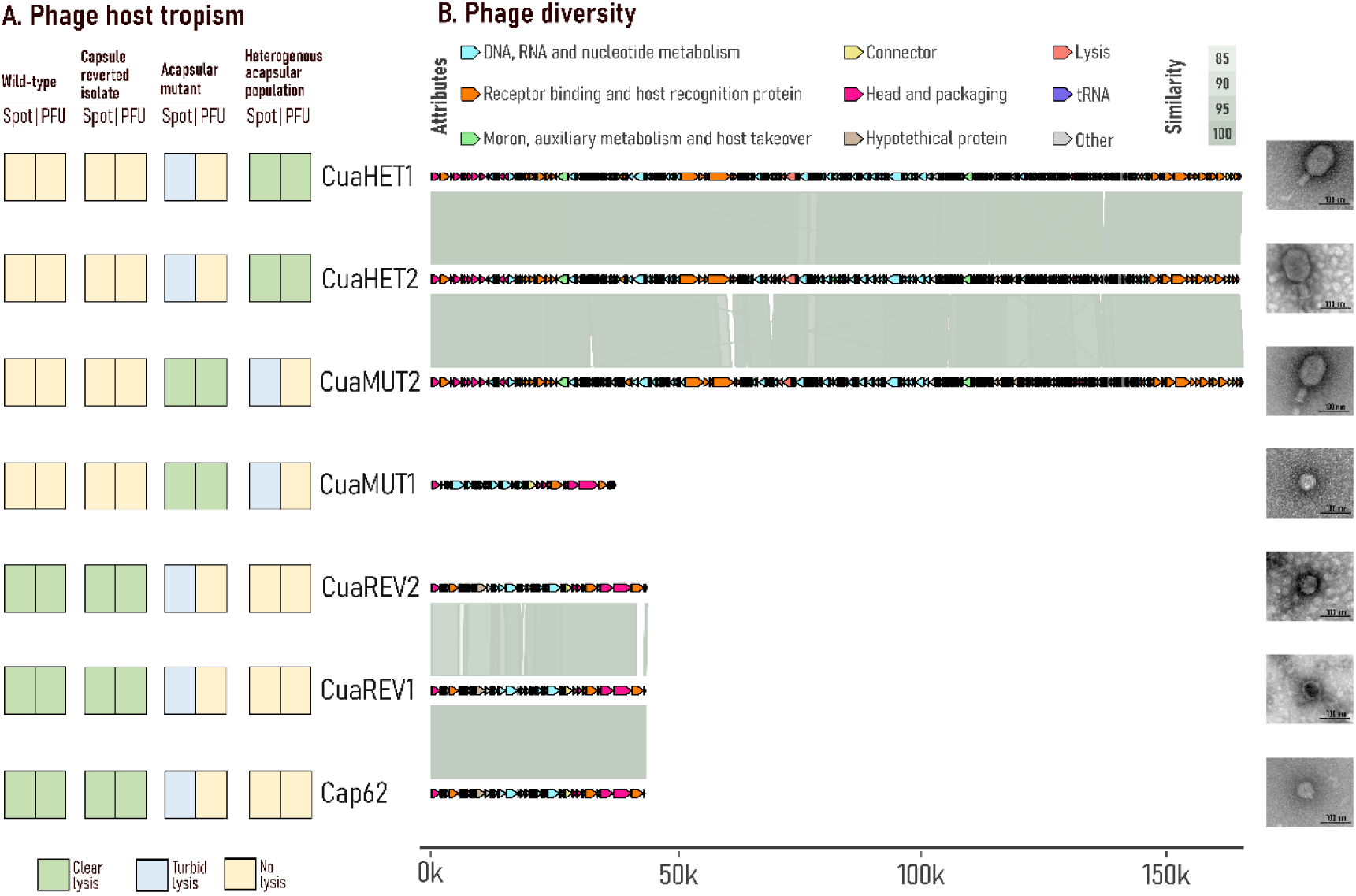
Phage host tropism and diversity. **A.** Phage host tropism assessed by the ability of phages to form spots or plaque forming units (PFU). Yellow boxes indicate no lysis, blue indicates turbid lysis, and green indicates clear lysis. **B.** Phage diversity analysis based on the intergenomic similarity and transmission electron microscopy. Arrows represent CDSs and are coloured based on functional groups. The grey scale represents the percentage of genomic similarity. The genomic scale is represented in base-pairs.

Full-genome sequencing revealed that phages CuaHET1, CuaHET2 and CuaMUT2 belonged to the *Jiaodovirus* genus (91.8 - 97.4 % intergenomic similarity). Phage CuaMUT1 belonged to the *Teetrevirus* genus. Finally, phages CuaREV1 and CuaREV2 belonged to the *Drulisvirus* genus, showing a high intergenomic similarity (85.1 - 99.9 %) with Cap62 (Figure 2B and Figure S1A). A virulent lifestyle was predicted for all phages (ranging from 96.2 to 100 %; Figure S1B). Depolymerase-related genes were just found in those phages belonging to the genus *Drulisvirus* and in phage CuaMUT1 (Figure S1C).

### Evaluation of phage cocktails to counteract phage resistance

To test the ability of these newly isolated phages to delay the emergence of phage-resistant bacteria, phage cocktails combining each individual isolated phage with phage Cap62 were tested to evaluate infectivity of WT culture in liquid media. The cocktails composed of CuaREV1 or CuaREV2 with Cap62 showed no delay in resistance, resulting in the same effect as that of Cap62 in monoinfection (*p-value > 0.1, paired t-test*). Interestingly, phage CuaMUT1 or CuaMUT2, together with Cap62, showed a slight delay in the emergence of resistance (*p-value < 0.05, paired t-test*). However, the greatest effect was shown by the combination of CuaHET1 or CuaHET2 with Cap62 (*p-value < 0.05, paired t-test*), where the resistance was delayed by more than 10 hours (Figure 3).

**Figure 3.**
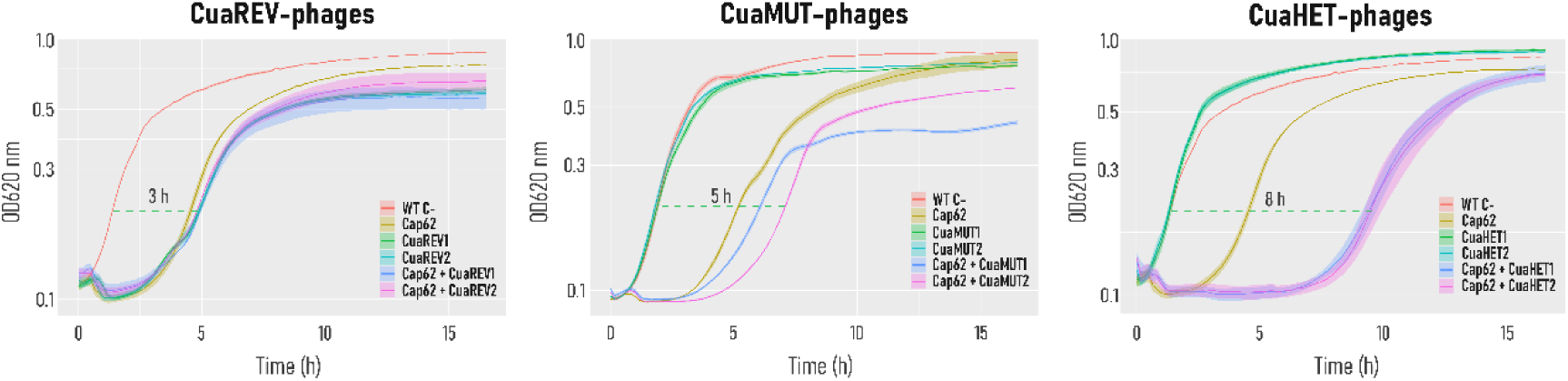
Efficacy of phage cocktails in delaying resistance emergence. Bacterial growth curves were measured by optical density at 620 nm over 16 hours. WT C-represents bacterial growth without phage. Other conditions represent bacterial growth with either a single phage or a cocktail of two phages. The shaded areas denote the standard error of the mean for three replicates.

### The delay in resistance is due to the complementary cell tropism by phages

To better understand the synergistic effect in delaying resistance, we focused on the phage cocktail encompassing Cap62 and CuaHET1, which proved to be one of the most effective combinations. To achieve this, the WT culture was passed through a flow cytometer, and individual cells were dispensed into wells containing different treatments: LB medium, phage Cap62, phage CuaH1, or a combination of both (phage cocktail). In the LB control wells, all cells were expected to grow, as no phage was present. In contrast, growth in wells containing phages was expected only from resistant cells. After 48 hours of incubation, we counted the number of wells in which bacteria had survived and exhibited growth. This allowed to estimate the proportion of cells in each culture that were sensitive to phage Cap62 and CuaHET1 (Figure 4A). The WT culture was predominantly composed of cells sensitive to Cap62, while 78% of the population was resistant to the CuaHET1 phage (Figure 4B). This confirmed that the WT was highly susceptible to Cap62, with Cap62-resistant variants present at levels below the detection threshold. Notably, 22% of the WT population remained sensitive to CuaHET1. Finally, no growth was observed in wells treated with the phage cocktail, further indicating that cells resistant to both phages are present at undetectably low frequencies. To assess whether resistance to Cap62 affects single-cell phage sensitivity, this assay was repeated using the heterogeneous Cap62-resistant population (Figure 4B). In this culture, the proportion of Cap62-resistant cells increased markedly to 92%, accompanied by a corresponding rise in sensitivity to CuaH1. This indicated that the acapsular phenotype promoted CuaHET1 infection. Conversely, analysis of a CuaHET1-resistant culture revealed a higher proportion of CuaHET1-resistant cells compared to the WT, while sensitivity to Cap62 was maintained. This reciprocal sensitivity likely underlies the delayed emergence of resistance observed when both phages were applied in combination (Figure 4C).

**Figure 4.**
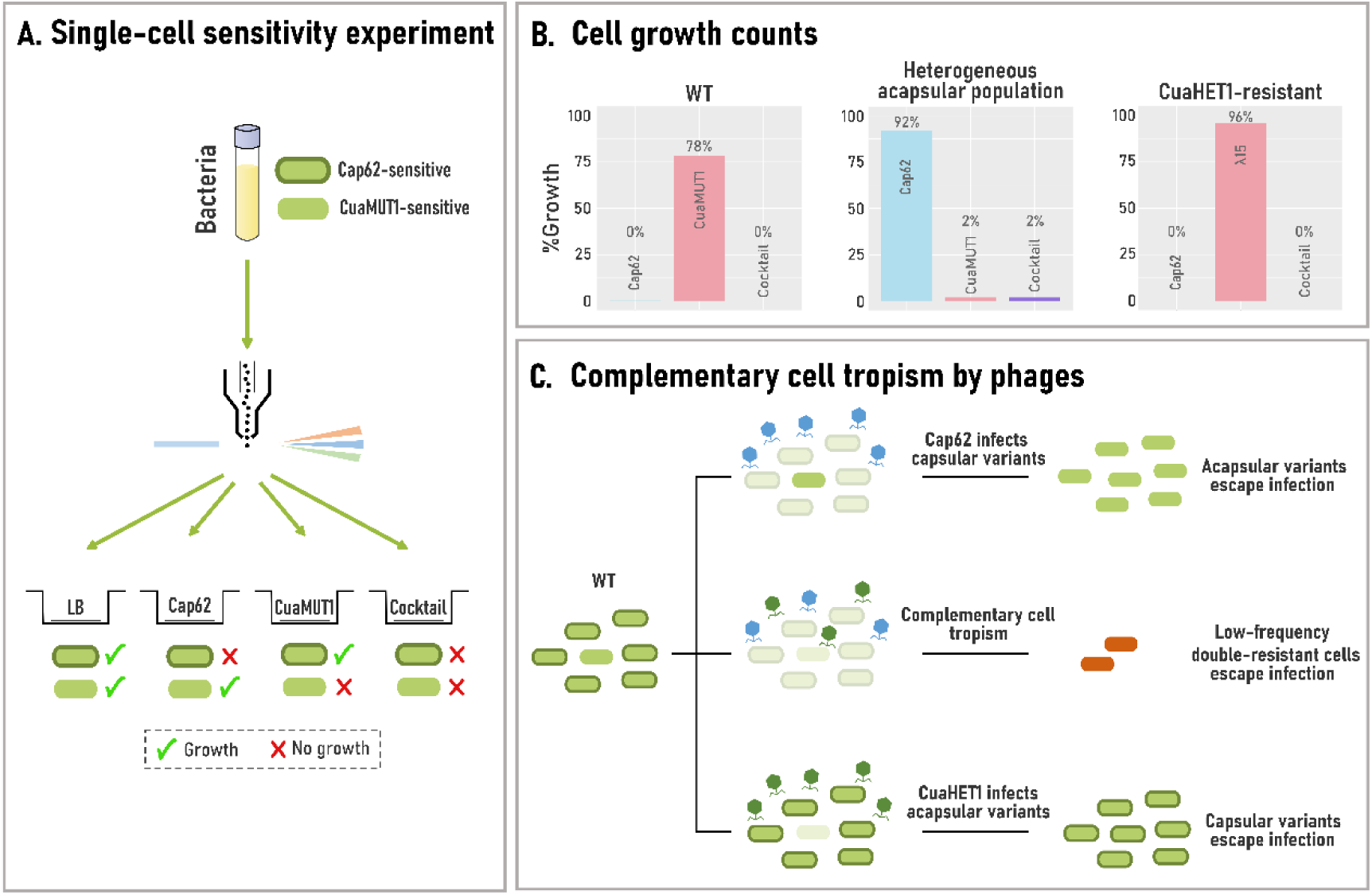
Single-cell sensitivity to phages. **A.** Schematic representation of the single-cell sensitivity experiment. Cells from bacterial cultures (WT, heterogeneous acapsular population, and CuaHET1-resistant) were sorted into individual wells containing different treatments (LB, Cap62, CuaHET1, or phage cocktail). Bacterial cells are shown in green, with a dark green outline indicating the presence of capsule. Capsulated cells are sensitive to Cap62, while acapsular cells are sensitive to CuaHET1. A green tick indicates bacterial growth, while a red cross indicates no growth. **B.** Percentage of wells with bacterial growth under each treatment, relative to the LB condition (100%). **C.** Schematic representation of the synergistic effect in delaying resistance emergence, driven by the complementary cell tropism of phages Cap62 (blue) and CuaHET1 (green).

### Cap62 selective pressure prevents capsule reversion and sustains cocktail synergy

The synergistic effect of the phage cocktail was driven by the differential lytic activity of both phages, Cap62 targeting capsular variants, and phage CuaHET1 acapsular ones. However, as previously shown, the resistant culture represented a heterogeneous population of acapsular variants. In the absence of Cap62-mediated selective pressure, some of these variants reverted to capsule production (e.g., the capsule-reverted isolate), while other remained acapsular (e.g., the acapsular mutant). This capacity for capsule reversion could undermine the long-term efficacy of the phage cocktail. To determine whether the presence of Cap62 was required to maintain CuaHET1 susceptibility, we isolated individual cells from the heterogeneous acapsular resistant population and allowed them to grow in wells containing either LB or phage Cap62 (Figure 5A). Following growth, cultures were subjected to single-cell sorting to assess their sensitivity to CuaHET1, and examined under the microscope to evaluate the presence of capsule (Figure 5B). The results indicated that acapsular cells grown in LB restored capsule production, leading to increased resistance to CuaHET1 (44% growth) compared to the original resistant population. In contrast, cells grown in the presence of Cap62 maintained the acapsular state and exhibited higher susceptibility to CuaREV1 (18% growth). Upon a subsequent passage, this trend was even more pronounced: cells grown in LB exhibited further increased resistance to CuaHET1 (60% growth), whereas those maintained under Cap62 pressure remained fully sensitive (0% growth). These findings suggest that Cap62 is essential for preserving the acapsular phenotype, thereby sustaining CuaHET1 efficacy within the cocktail.

**Figure 5.**
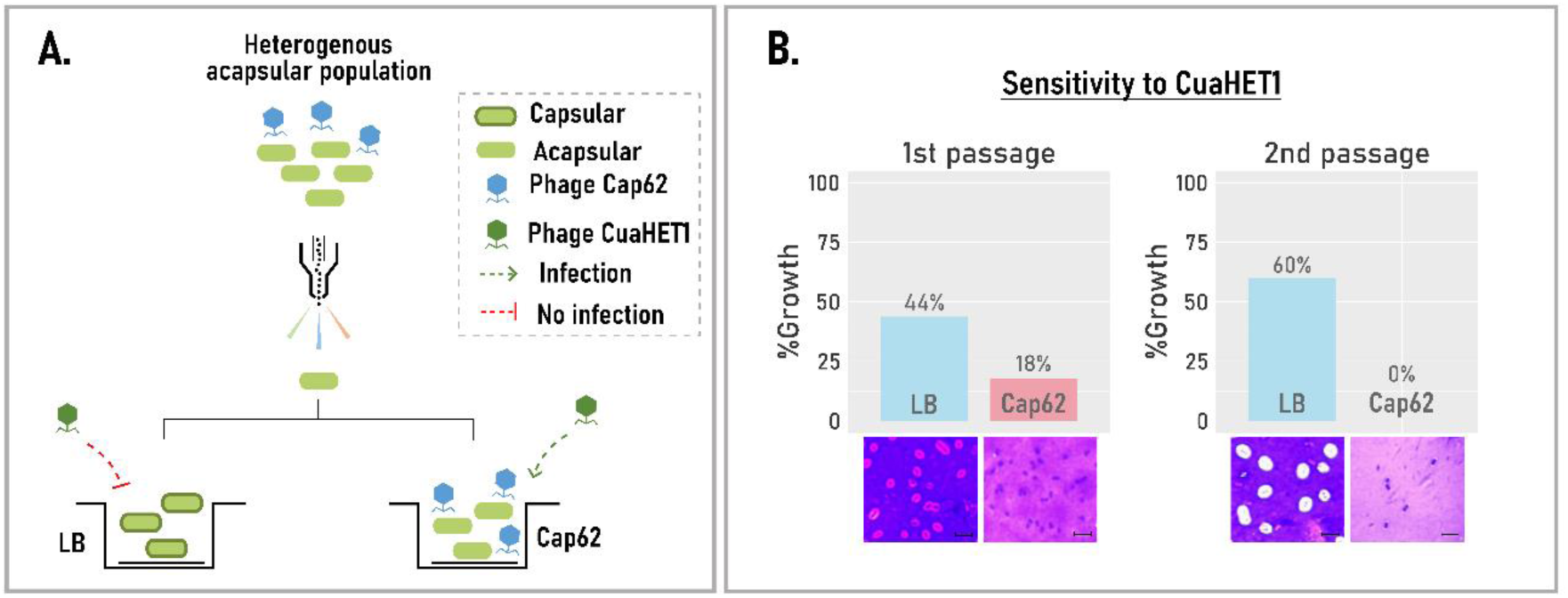
Influence of phage Cap62 on capsule dynamics and susceptibility to CuaHET1. **A.** Schematic representation of the tested hypothesis. Cells from the heterogeneous acapsular population grown in LB restore capsule and become resistant to phage CuaHET1 (red dotted line), whereas those grown in the presence of Cap62 remain acapsular and susceptible to phage CuaHET1 (green dotted line). **B.** Sensitivity of single-cell acapsular derived cultures grown in LB (blue) or in the presence of Cap62 (salmon) to phage CuaHET1, measured as the percentage of wells with bacterial growth (%Growth) over two consecutive single-cell passages. Microscopy images show that cells grown in LB restore capsule production, while cells grown in Cap62 retain the acapsular phenotype. The scale bars represent 5 µm.

## DISCUSSION

The emergence of phage-resistant bacteria is a major challenge for phage therapy (5). While phage cocktails have shown efficacy in delaying resistance, their effectiveness depends on the ability to target the full spectrum of resistant variants (36). Most studies isolated and characterized resistance based on clonal cultures (32). However, phage-resistant populations often display substantial genetic and phenotypic diversity, which cannot be only represented by a single clone (37). In this study we demonstrated that distinct subpopulations within phage-resistant *K. pneumoniae* cultures displayed differential susceptibility to phages, highlighting how population heterogeneity can shape phage-host interactions and influence cocktail efficacy.

Previous studies have shown that capsule-deficient mutants facilitate the isolation of phages that specifically target acapsular variants, while capsulated strains typically yield capsule-dependent phages (14, 19, 38). In some cases, multiple acapsular mutants were isolated to cover the full resistance spectrum (32, 39, 40). However, this approach is not always feasible, especially when the resistant population includes transient acapsular variants (35). In such cases, applying a selective pressure, such as a capsule-targeting phage, can help to stabilize the acapsular state and prevent capsule reversion, thereby enhancing the isolation of phages that target these unstable phenotypes. Phages capable of targeting a broader range of resistant variants are generally more effective at delaying resistance emergence (19, 31, 32). This effect is often attributed to complementary host tropism, where each phage targets a distinct subset of the bacterial population (36, 41). Therefore, combinations of phages with overlapping host ranges, such as Cap62 and CuaREV-like phages, may be less effective due to functional redundancy. In contrast, combinations that include phages targeting distinct bacterial subpopulations, such as capsulated and acapsulated variants, are more likely to exert broad selective pressure and reduce the chances of resistant escape.

Isolating phages from heterogeneous resistant populations may increase the chance of selecting phages with broader host ranges, whereas relying on clonal cultures like acapsular mutants may limit this potential (42). However, in this study, the superior efficacy of CuaHET-like phages compared to CuaMUT-like phages does not appear to be due to a broader host range, as CuaHET-like phages were unable to effectively infect the acapsular mutant culture (43). An alternative explanation is the presence of a dominant subpopulation within the heterogeneous resistant culture. Phages able to target this subpopulation may exert stronger selective pressure, resulting in enhanced suppression of resistance compared to phages targeting less abundant subpopulations (44). Although capsule biosynthesis in *K. pneumoniae* is primarily governed by the K-locus, additional genes can modulate capsule quantity and architecture, potentially affecting access to alternative receptors such as LPS or OMPs (21). Moreover, LPS and OMP may also vary across cells within the population (22, 23), adding a layer of heterogeneity that shapes population sensitivity to these phages. As a result, the acapsular phenotypes of a mutant clone and a reversible acapsular variant may result in different susceptibility to non-capsule-dependent phages, as for the case of CuaHET-like and CuaMUT-like phages. Identifying the receptors used by these phages would clarify these differences, enabling a more targeted selection of complementary phages for improved cocktail design.

Our previous findings demonstrated that, although acapsular mutants are present in the resistant population, mutation-driven acapsularity is not the predominant resistance mechanism (35). Instead, resistance mostly arises from transient acapsular states, evidenced by the downregulation of capsule biosynthesis genes and the rapid restoration of capsule expression observed in single-cell experiments. This reversible phenotypic switch may result from mechanisms such as phase variation or stochastic fluctuations in gene expression, which facilitate adaptation to challenging environments (45). While spontaneous mutations in bacteria occur at a relatively low frequency, around 10⁻¹⁰ mutations per nucleotide per generation (46, 47), phase variation can occur at rates up to three orders of magnitude higher (24). Stochastic gene expression changes are even more frequent, generating phenotypic variability on much shorter timescales (26, 27, 48, 49). Consequently, the superior ability of CuaHET-like phages to delay resistance may stem from their capacity to infect non-mutational acapsular variants, which likely emerge more frequently than acapsular mutants in the population.

Single-cell experiments underscored the dynamic interplay between phage pressure and bacterial phenotype. The efficacy of CuaHET-like phages seems to be tightly linked to the selective pressure exerted by a capsule-targeting phage, necessary to maintain the acapsular state. Once this pressure is removed, capsule expression can be rapidly restored, reducing phage susceptibility. These observations support the idea that simultaneous administration of complementary phages may be more effective than sequential treatments in capsular bacteria (50), as it maintains selective pressure and prevents phenotypic reversion that could compromise cocktail efficacy. Although *in vivo* studies have shown that acapsular variants are more susceptible to the immune system and antibiotics (28), the contribution of CuaHET-like phages may still be critical in specific clinical contexts. In particular, in immunocompromised patients or in situations where antibiotic options are limited or no longer viable, phage cocktails targeting both capsular and acapsular subpopulations may provide a valuable control strategy.

In conclusion, our findings highlight the importance of considering phenotypic diversity within bacterial populations when designing phage therapies. The effectiveness of phage cocktails depends not only on combining phages targeting distinct receptors but also on understanding how selective pressures shape population phenotype and influence phage susceptibility. These results underscore the limitations of conventional phage selection strategies and support population-level approaches to improve the efficacy of phage therapy.

## METHODS

### Bacterial strains and phage Cap62

The reference *K. pneumoniae* strain used in this study corresponds to the capsular type 1 strain (referred to as WT), obtained from the Statens Serum Institut (Copenhagen, Denmark). Phage Cap62 and the three derived phage-resistant cultures were described previously (35). These cultures include: the heterogeneous acapsular population composed primarily of reversible acapsular variants, along with a minority of stable acapsular mutants; an acapsular mutant with a stable phenotype; and a capsule-reverted isolate that harboured no detectable mutations and restored capsule production through changes in gene expression All bacterial cultures were grown in lysogenic broth (LB) supplemented with 3.78 mM CaCl_2_ to promote phage adsorption at 37 °C with shaking.

### Isolation and purification of phages

Wastewater samples from the metropolitan area of Valencia (Spain) were centrifuged at 4 °C, 4000 × g for 10 min to pellet dust and large particles. The supernatant was filtered through 0.22 µm filters and 800 µL was mixed with 200 µL of a Cap62-resistant culture on stationary phase. The mixture was incubated at room temperature for 15 min, and then mixed with 3.5 mL of semi-solid LB agar and poured onto LB agar plates. After an overnight incubation at 37 °C, lytic plaques were selected for plaque-to-plaque purification with a minimum of three passages. For each passage, plaques were resuspended with LB, centrifuged to pellet bacteria, and the supernatants were diluted and plated for the next passage. The supernatant from the final purification passage was amplified in LB + CaCl_2_ using 10^6^ – 10^7^ colony-forming units (CFU)/mL and an initial phage concentration ranging between 10^6^ – 10^7^ PFU/mL. After approximately 3 h of incubation at 37 °C with shaking, samples were centrifuged at 18000 ×g for 3 min to pellet bacteria, and the supernatant was filtered through 0.22 µm filters. Finally, samples were aliquoted and stored at -70 °C.

### Phage infectivity via spot assay and plaque-forming unit analysis

To assess the efficiency of phages in infecting the phage resistant cultures, spot and plaque assays were done to test the ability of phages to form PFUs. For spot-tests, 200 µL of a stationary-phase culture containing approximately 10^9^ CFU was mixed with 3.5 mL of semi-solid agar LB and plated onto an LB agar plate. After drying, 1 µL of phage was spotted at a concentration of approximately 10^8^ PFU/mL, followed by an overnight (ON) incubation at 37 °C. Additionally, ten-fold serial dilutions were routinely spotted. To evaluate the ability of phages to form PFU in the resistant cultures, serial dilutions of the phage were prepared, and 10 µL of each dilution was plated with bacteria in semi-solid LB agar, as previously described. After ON incubation at 37 °C, the presence of PFUs on the agar plates was assessed. Each experiment was evaluated in at least three independent experiments, and the results were scored as clear lysis, turbid lysis or no lysis.

### Bacterial Growth Curve Assay to Evaluate Phage Cocktail Efficacy

To test the ability of cocktails to delay the emergence of resistant bacteria, the newly isolated phages were individually combined with Cap62 in liquid infections. Approximately 5 × 10^6^ CFU of the WT culture was mixed with phages at a total multiplicity of infection (MOI) of 0.5 in a final volume of 150 µL. In the combined treatments, the amount of each phage was half that used in the mono-infection. To monitor phage-induced lysis and the growth of resistant bacteria, we measured OD_620_ nm over 16 h with continuous shaking at 37 °C, using 96-well plates and a plate reader (Multiskan FC). Experiments were performed in triplicate. We determined the time of resistance emergence as the difference between the maximum slope of the uninfected and infected cultures. The slope was calculated 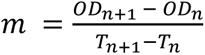, where *OD* represents the OD_620_ nm, and *T* is the time.

### Capsule staining

To determine the presence of the capsule in bacterial cultures, contrast staining with crystal violet and nigrosine was performed as described in (51). Briefly, liquid cultures with a minimum recommended concentration of 5 × 10^9^ CFU/mL were fixed for 20 min with 2.5% formaldehyde in the presence of 100 mM lysine. The fixative agent was removed by centrifugation and washed with 1× phosphate-buffered saline (PBS). 25 – 40 µL of the fixed culture was mixed with a small drop of nigrosine 10% in a glass slide and smeared along the slide until air dried. 1% crystal violet was gently poured to cover the slide and incubated for 5 min at room temperature. The preparation was carefully rinsed with distilled water, air dried and examined under a light microscope.

### Single-cell sensitivity to phages

The WT culture was grown in LB medium alone, and in the presence of phage (10^8^ PFU/mL) to generate the phage-resistant cultures. To remove the broth culture and partially eliminate the surrounding phage, cultures were washed five times with 1× PBS. These were then resuspended in PBS to a final concentration of 10^5^ CFU/mL. The cultures were subsequently passed through a cytometer (BD FACSAria™ Fusion) under biosafety level-2 conditions to isolate individual cells into 96-well plates containing different treatments: LB, phage Cap62, phage CuaH1, or a cocktail of both phages. Phage concentrations were maintained extremely high (10^8^ – 10^9^ PFU/mL) to ensure that only resistant bacteria could grow. Plates were incubated for 48 h at 37 °C, and we counted the number of wells in which bacteria grew. The number of wells with bacterial growth in LB medium was considered as the maximum growth (100%). The proportion of wells with bacterial growth in the presence of phage was then normalized to this maximum growth to determine the relative resistance of the bacteria to the phages.

To analyze the effect of phage selective pressure on the maintenance of acapsularity, we performed a first passage, isolating single cells from the heterogeneous acapsular population into wells containing LB or Cap62 (10^8^ – 10^9^ PFU/mL). After 48 h incubation at 37 °C, cultures were subjected to a second passage via single-cell sorting, in which individual cells were again isolated in LB or Cap62. Both passages were subjected to cell sorting, and individual cells were exposed to LB and Cap62 (10^8^ – 10^9^ PFU/mL) treatments. Plates were incubated, and and the number of wells with bacterial growth was counted as previously described.

### Transmission electron microscopy of phages

Phage lysates were filtered through 0.22 µm and concentrated by centrifugation at 80000 × g, 4 °C, 2 h to obtain a minimum titer of 10^10^ PFU/mL. The pellet was resuspended in SM buffer and a small volume placed on a carbon-coated Formvar supported by a 300 mesh copper grid. Preparations were air dried for 30 min and any excess of liquid was removed with filter paper. The samples were then negatively stained with 2% phosphotungstic acid and observed using a Jeol JEM-1010 electron microscope.

### Phage sequencing and genomic analysis

Phage lysates were filtered through 0.22 µm and concentrated using the Concentrating Pipette Select (Innovaprep) to reach a minimum titer of 10^10^ PFU/mL. 180 µL of the concentrated lysate was used to extract DNA from viral capsids using an automated protocol with the Maxwell RSC Instrument (Promega). DNA libraries were generated using the Illumina Nextera XT DNA kit (2 × 150 bp or 2 × 250 bp paired-end reads), and the Illumina MiSeq platform was used to generate the sequencing reads with the MiSeq Reagent Kit v2. Read quality was checked using FastQC (52) and genomes were assembled *de novo* using SPAdes v3.15.4 (53) in a single contig. Phage genomes were annotated using Pharokka (54). Genes were classified into functional groups assigned by PHROGs and a comparative plot of the genomes was made using gggnomes (55). BACPHLIP (56) was used to predict the lifestyle of phages. And VIRIDIC (57) was used to estimate the intergenomic similarity between phages. To predict depolymerase activity in phage coding sequences (CDS), three tools, DepoScope (58), PhageDPO (59) and DePP (60) were used. Proteins were considered depolymerases only if all three predictors returned a score ≥ 0.90.

## Author contributions

Conceptualization, L.M-Q., R.S. and P.D-C.; investigation, L M-Q.; writing— original draft, L.M-Q.; writing—review & editing, R.S. and P.D-C.; funding acquisition, P.D-C.

## Data availability

Unrestricted access to all data: P.D.-C.

