## Supplementary material for "Phenotypic Heterogeneity Shapes Phage Resistance and Cocktail Efficacy in *Klebsiella pneumoniae*": Figure S1

A. Pairwise intergenomic similarity (VIRIDIC)

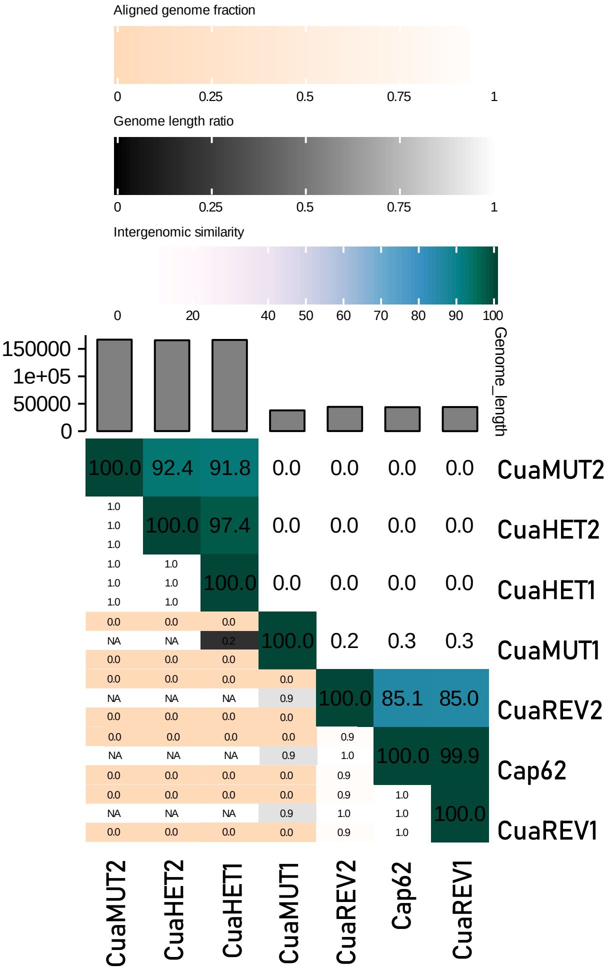

### B. Predicted phage lifestyle (BACPHLIP)

| Phage | Virulent | Temperate |
|---|---|---|
| **Cap62** | 1.0 | 0.0 |
| **CuaREV1** | 1.0 | 0.0 |
| **CuaREV2** | 0.9752481267495525 | 0.024751873250447337 |
| **CuaMUT1** | 0.9981617952123443 | 0.001838204787655567 |
| **CuaMUT2** | 0.9745052770448549 | 0.02549472295514512 |
| **CuaHET1** | 0.9620052770448548 | 0.037994722955145124 |
| **CuaHET2** | 0.9620052770448548 | 0.037994722955145124 |

### **C.** Depolymerase prediction

| Phage | CDS | Name | DepoScope | PhageDPO | DePP |
|---|---|---|---|---|---|
| **Cap62** | 007 | Tail fiber protein | 1.00 | 0.99 | 0.94 |
| | 056 | Tail protein | 1.00 | 0.91 | 0.98 |
| | 060 | Tail fiber protein | 1.00 | 1.00 | 0.96 |
| **CuaREV1** | 007 | Tail fiber protein | 1.00 | 0.99 | 0.95 |
| | 056 | Tail protein | 1.00 | 0.91 | 0.97 |
| | 060 | Tail fiber protein | 1.00 | 1.00 | 0.95 |
| **CuaREV2** | 007 | Tail fiber protein | 1.00 | 0.97 | 0.96 |
| | 056 | Tail protein | 1.00 | 0.93 | 0.96 |
| | 060 | Tail fiber protein | 1.00 | 0.95 | 0.97 |
| **CuaMUT1** | 046 | Tail protein | 1.00 | 0.97 | 0.96 |
| **CuaMUT2** | nd | nd | - | - | - |
| **CuaHET1** | nd | nd | - | - | - |
| **CuaHET2** | nd | nd | - | - | - |

Phage coding sequences with predicted score ≥ 0.90 by all three depolymerase predictors are shown.

nd: No coding sequences were predicted ass depolymeraes by all three predictors.
